# Virtual brain simulations reveal network-specific parameters in neurodegenerative dementias

**DOI:** 10.1101/2023.03.10.532087

**Authors:** Anita Monteverdi, Fulvia Palesi, Michael Schirner, Francesca Argentino, Mariateresa Merante, Alberto Redolfi, Francesca Conca, Laura Mazzocchi, Stefano F. Cappa, Matteo Cotta Ramusino, Alfredo Costa, Anna Pichiecchio, Lisa M. Farina, Viktor Jirsa, Petra Ritter, Claudia A.M. Gandini Wheeler-Kingshott, Egidio D’Angelo

## Abstract

**Introduction:** Neural circuit alterations lay at the core of brain physiopathology, and yet are hard to unveil in living subjects. Virtual brain modelling (TVB), by exploiting structural and functional MRI, yields mesoscopic parameters of connectivity and synaptic transmission.

**Methods:** We used TVB to simulate brain networks, which are key for human brain function, in Alzheimer’s disease (AD) and Frontotemporal Dementia (FTD) patients, whose connectivity and synaptic parameters remain largely unknown; we then compared them to healthy controls, to reveal novel in vivo pathological hallmarks.

**Results:** The pattern of simulated parameter differed between AD and FTD, shedding light on disease-specific alterations in brain networks. Individual subjects displayed subtle differences in network parameter patterns that significantly correlated with their individual neuropsychological, clinical, and pharmacological profiles.

**Discussion:** These TVB simulations, by informing about a new personalized set of networks parameters, open new perspectives for understanding dementias mechanisms and design personalized therapeutic approaches.

## INTRODUCTION

The advent of advanced in human in vivo recordings of brain signals from e.g., magnetic resonance imaging (MRI), has led to the identification of brain networks that subtend specific functions [1]. The structural and/or functional alteration of such networks eventually leads to the clinical manifestation of neurological diseases. In parallel, mathematical modelling of cellular and microcircuit functions are emerging, providing tools to link the micro- to the meso- and the macro-scale properties of brain signals [2].

Neurodegenerative dementias include several neuropathological forms, primarily Alzheimer disease (AD) and frontotemporal dementia (FTD). Post-mortem histology and in vivo functional MRI (fMRI) studies have suggested a differential engagement of various brain networks in these diseases. However, a comprehensive assessment of functional connectivity (FC) changes in multiple networks in vivo to compare dementias subtypes has been rarely performed [3], in favor of investigating specific networks, in particular the default mode network (DMN) specifically in AD [4]. Increasing evidence underlines the need to expand the investigation beyond the DMN, considering that widespread increases and decreases in structural and functional connectivity have been observed in different brain areas of AD patients [5]. Moreover, the development of in vivo imaging biomarkers of brain function becomes necessary to achieve efficient tailored diagnosis and personalized treatment, especially in less frequent and more heterogeneous conditions, such as atypical forms of AD or FTD variants [6].

Advanced recording techniques, such as MRI and/or high-density Electroencephalography (hd-EEG), are mostly used to study structural and functional brain networks properties and their changes in pathological conditions, but they provide little information about cellular properties such as spatio-temporal dynamics of cellular communication, neuronal firing integrity or synaptic transmission. Therefore, very little is known about the cellular and synaptic changes typical of different diseases, and even more so about whether changes that have cascaded from cells to networks are specific to individual patients.

Recent advances in multiscale brain modelling offer promising tools to study the whole brain temporal dynamics, integrating macroscopic information from structural and functional MRI with mathematical mesoscale representations of the underlying ensemble properties of cells and microcircuits. In particular, The Virtual Brain (TVB) modelling workflow allows the non-invasive investigation of brain features, such as network connectivity strength and excitatory/inhibitory (E/I) balance [2,5], which are relevant to brain disease and can be determined for each patient. The E/I balance, in turn, can be extracted at whole brain level or for specific brain networks from parameters measuring excitatory coupling, inhibitory coupling, and recurrent excitation inside network nodes [7]. Importantly, all neurological conditions involve changes at multiple scales and can gain from the use of TVB for understanding the impact of cellular and microcircuit properties alterations on brain function. The promise for clinical use of TVB has been already suggested in epilepsy surgery [8], stroke [9], brain tumors [10], Multiple Sclerosis [11] and neurodegenerative conditions like dementia [12-15]. Interestingly, the central position of an E/I imbalance in the cascade of pathophysiological events in AD is increasingly recognized [16]. However, very little is known on how such network neurophysiology acts in concert with structural and functional connectivity alterations to determine cognitive decline. Retrieving E/I information, even if summarized in mesoscale network parameters, is extremely important, as it will provide new insights in neurodegenerative mechanisms of disease that will eventually impact on finding effective treatments.

In this work, we applied TVB to enable the non-invasive investigation of connectivity strength and E/I balance in a heterogeneous cohort of dementia patients, including typical and atypical AD and FTD variants. We explored the relationship between neurophysiological parameters provided by TVB in multiple brain networks and neuropsychological scores recorded during patient examinations. TVB parameters differentiated AD from FTD and proved to be sensitive to profiles of cognitive performance and ongoing pharmacological treatment. In aggregate, this study shows how TVB analysis can be used to provide personalized fingerprints of dementia patients, opening new perspectives for differential diagnosis and for tailoring pharmacological and interventional workflows.

## MATERIALS AND METHODS

### Subjects

Twenty-three patients affected by neurodegenerative diseases were recruited at the IRCCS Mondino Foundation. The study was approved by the local Ethical Committee and carried out in accordance with the Declaration of Helsinki. Written informed consent was obtained from all subjects. The protocol was approved by the local ethical committee of the IRCCS Mondino Foundation. Patients underwent a complete diagnostic workup including clinical and neuropsychological assessment (see section below) MRI, and, when available, cerebrospinal fluid (CSF) biomarkers (amyloid-β and τ protein) assessment following the harmonized protocol of the RIN network (Italian Network of the Institutes (IRCCS) of Neuroscience and Neurorehabilitation) [17]. Subjects were classified into two main groups: 16 AD patients (13 females, 70 ± 8 years) and 7 FTD patients (1 female, 69 ± 5 years), further classified into distinct phenotypes. In particular, AD patients were additionally classified into: typical AD (10 subjects, [18]); AD logopenic variant (2 subjects, [18]); AD frontal variant (ADfv, 1 subject, [18]); AD posterior cortical atrophy (ADpca, 1 subject,[18]). One patient was classified as having corticobasal syndrome (CBS, 1 subject, [19]) and one with Dementia with Lewy bodies (DLB, 1 subject,[20]). On the other hand, FTD patients were classified into: behavioral FTD (FTDbv, 5 subjects, [21]); Primary Progressive Aphasia non-fluent variant (PPAnf, 1 subject, [22]) and Primary Progressive Aphasia semantic variant (PPAsv, 1 subject, [22]). Pharmacological therapy was also recorded.

10 Healthy Controls (HC, 6 females, 67 ± 3 years) were enrolled on a voluntary basis as reference group. All HC underwent clinical assessment to exclude any cognitive impairment. For all subjects, exclusion criteria were: age>80 years, a diagnosis of significant medical, neurological and psychiatric disorder, pharmacologically treated delirium or hallucinations and secondary causes of cognitive decline (e.g. vascular metabolic, endocrine, toxic and iatrogenic). Suppl Table 1 shows demographic, clinical, and neuropsychological data.

### Neuropsychological assessment

All subjects underwent a neuropsychological examination based on a standardized battery of tests to assess their global cognitive status (Mini-Mental State Examination, MMSE) and different cognitive domains: memory (verbal: Rey’s Auditory Verbal Learning Test, RAVLT; visuo-spatial: Rey– Osterrieth complex figure recall), phonemic and semantic fluency, visuo-constructional abilities (Rey–Osterrieth complex figure copy), attention (Trial Making Test part A, TMT-A) and executive functions (Frontal Assessment Battery, FAB; Trial Making Test part B and B-A; Stroop colour-word test interference, time and errors; Raven’s Coloured Progressive Matrices, CPM47).

Raw scores were corrected for the effect of age, education, and gender according to the reference norms for the Italian population. Corrected scores were classified into five Equivalent Scores (ES), from 0 to 4, with an ES of 0 reflecting a pathological performance. Domain scores, calculated by averaging the ES of the single tests, were obtained for memory, language-fluency, visuo-constructional abilities, attention, and executive functions, respectively.

### MRI acquisitions

All subjects underwent MRI examination using a 3T Siemens Skyra scanner with a 32-channel head coil. The MRI protocol was harmonized within the RIN network including both diffusion weighted imaging (DWI) and resting-state fMRI (rs-fMRI) [17]. For DWI data a 2-shell standard single-shot echo-planar imaging sequence (EPI) (voxel size = 2.5 × 2.5 × 2.5 mm^3^, TR/TE = 8400/93 ms, two shells with 30 isotropically distributed diffusion-weighted directions, diffusion weightings of 1,000 and 2,000 s/mm^2^, 7 non-diffusion weighted b = 0 s/mm^2^ images (b_0_ images) interleaved with diffusion-weighted volumes) was implemented, and 3 non-diffusion weighted images with the reversed phase-encoding acquisition were additionally acquired for distortion correction. For the rs-fMRI data, GE-EPI sequence (voxel size = 3 × 3 × 3 mm^3^, TR/TE = 2400/30 ms, 200 volumes) was set. For anatomical reference, the protocol included a whole brain high-resolution 3D sagittal T1-weighted (3DT1) scan (TR/TE = 2300/2.96 ms, TI = 900 ms, flip angle = 9°, voxel size = 1 × 1 × 1 mm^3^).

### Preprocessing of DWI and fMRI data

Preprocessing of diffusion and fMRI data was performed according to [14]. Briefly, DWI data were denoised, and corrected for motion and eddy currents distortions, then white matter, gray matter (GM), subcortical GM and CSF were segmented from the co-registered 3DT1 volume. 30 million streamlines whole-brain anatomically constrained tractography [23] was performed within MRtrix3, estimating fibers orientation distribution with multi-shell multi-tissue constrained spherical deconvolution (CSD) and using probabilistic streamline tractography [24]. fMRI preprocessing included denoising, slice-timing correction, realignment, co-registration to the 3DT1 volume, polynomial detrending, nuisance regression of 24 motion parameters [25] and CSF temporal signal [26], and temporal band-pass filtering (0.008-0.09 Hz).

### Structural and functional connectivity

An ad-hoc anatomical atlas in MNI (Montreal Neurological Institute) space was created combining 93 cerebral (AAL) (including cortical/subcortical structures) and 33 cerebellar (SUIT) labels [27]. We then performed a mapping between our ad-hoc atlas and the Buckner and Yeo [28,29] cerebral and cerebellar functional atlases to select the grey matter anatomical nodes of six networks known to support specific functions: i) integrative networks: default mode network (DMN), frontoparietal network (FPN), limbic network (LN), attention network (AN); ii) motor and sensory networks: visual network (VN), somatomotor network (SMN) (Fig. 1). For each subject, the grey matter parcellation of our combined anatomical atlas was applied to the whole-brain tractography to extract a whole-brain structural connectivity (SC) matrix, with the normalized number of streamlines as edges and cortical/subcortical/cerebellar areas as nodes. The subset of nodes defining each network and their connections were extracted from whole-brain SC obtaining specific network SC matrices, used as input to TVB (as detailed below). In addition, both static and dynamic experimental FC (expFC and expFCD, respectively) were reconstructed from rs-fMRI data for each of the six brain networks, to capture not only synchronous fluctuations of BOLD signals but also their spatiotemporal-dynamics during resting-state [30]. The expFC matrix was created by extracting the time-course of BOLD signals for each node and computing the Pearson’s correlation coefficient (PCC) of the time-course of pairs of atlas-defined brain regions. Matrix elements were converted with a Fisher’s z transformation and thresholded at 0.1206 [31]. To obtain expFCD, expFC was first computed over a sliding window of 40 seconds (expFCsw), shifted incrementally by 1 repetition time, which for our data it means to have 178 expFCsw [32]. Then, expFCD was calculated as a time-versus-time matrix, containing the Pearson correlation between each expFCsw and all expFCsw, centered at all other time points along the total acquisition window, quantifying, therefore, time-evolving dynamics.

**Figure 1.**
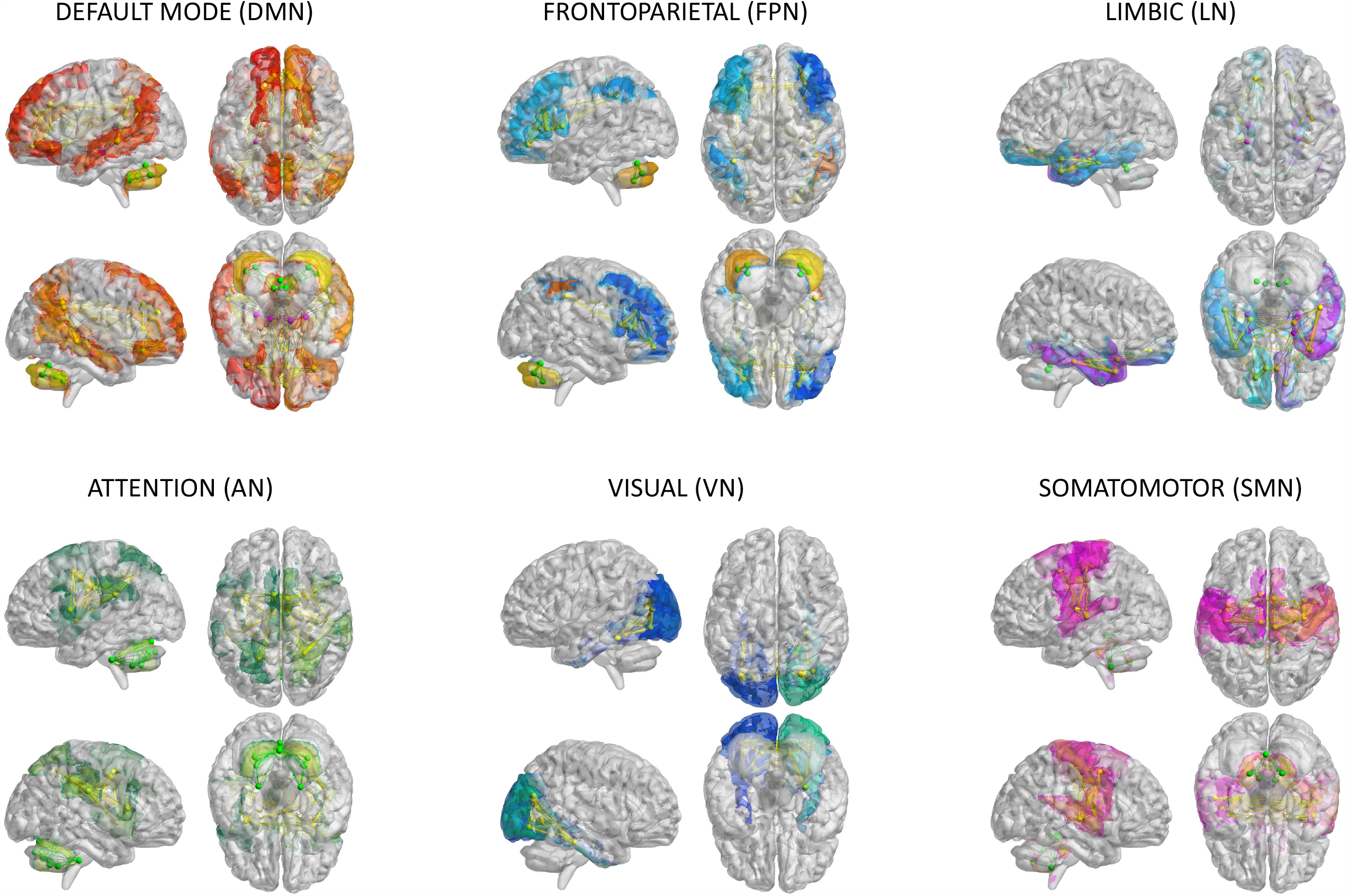
Brain networks. The six networks considered for modelling brain dynamics with the virtual brain (TVB): default-mode (DMN), frontoparietal (FPN), limbic (LN), attention (AN), visual (VN), somatomotor (SMN) network. These networks were defined according to Buckner and Yeo atlases and extracted from whole-brain structural connectivity matrices of each subject, choosing a subset of nodes and connections from the whole brain parcellation. Nodes and edges considered for each network are differently colored.

### Virtual brain modelling

The TVB workflow (reported in [14] for the whole brain) was applied to each one of the six selected brain networks (Fig. 2). The Wong-Wang neural mass model [7] (Suppl Fig.1), implemented with an optimized C code [33], was chosen to simulate local microcircuits activity, resulting from two populations of interconnected excitatory and inhibitory neurons coupled through NMDA and GABA receptor types. In our TVB simulations, this neural mass model was associated to each node of the network, while the SC matrix was used for the nodes interconnection. A set of parameters had to be tuned globally for each network: the global coupling (G), which is a scaling factor that represents the connections strength, and three synaptic parameters, i.e. the excitatory (NMDA) synapses (J_NMDA_), the inhibitory (GABA) synapses (J_i_), and the recurrent excitation (w_+_). The neural activity simulated with TVB was fed into the Balloon-Windkessel hemodynamic model [34] to reconstruct resting-state BOLD fMRI time-courses over 8 min length and compute simulated FC (simFC) and FCD (simFCD). Parameters were adjusted iteratively using expFC and expFCD of each network as targets to optimize model fitness and the validity of the result was assessed by iterating the optimization using different initial conditions (Suppl Fig.2) [35]. For the simFC vs. expFC comparison, model parameters were tuned until the PCC between experimental and simulated data reached the highest value. For the simFCD vs. expFCD comparison, differences between experimental and simulated FCD were assessed using the Kolmogorov-Smirnov (KS) distance: lower KS values corresponded to a lower distance of frame-by-frame FCD properties, meaning that model and experimental matrices were closest to each other. Thus, to achieve the optimal TVB simulation it was necessary to find both the highest PCC and the lowest KS values. To this aim, an overall cost function was defined as (1 - PCC) + KS and lowest cost function values implied the best fit both to static and dynamic functional data [36].

**Figure 2.**
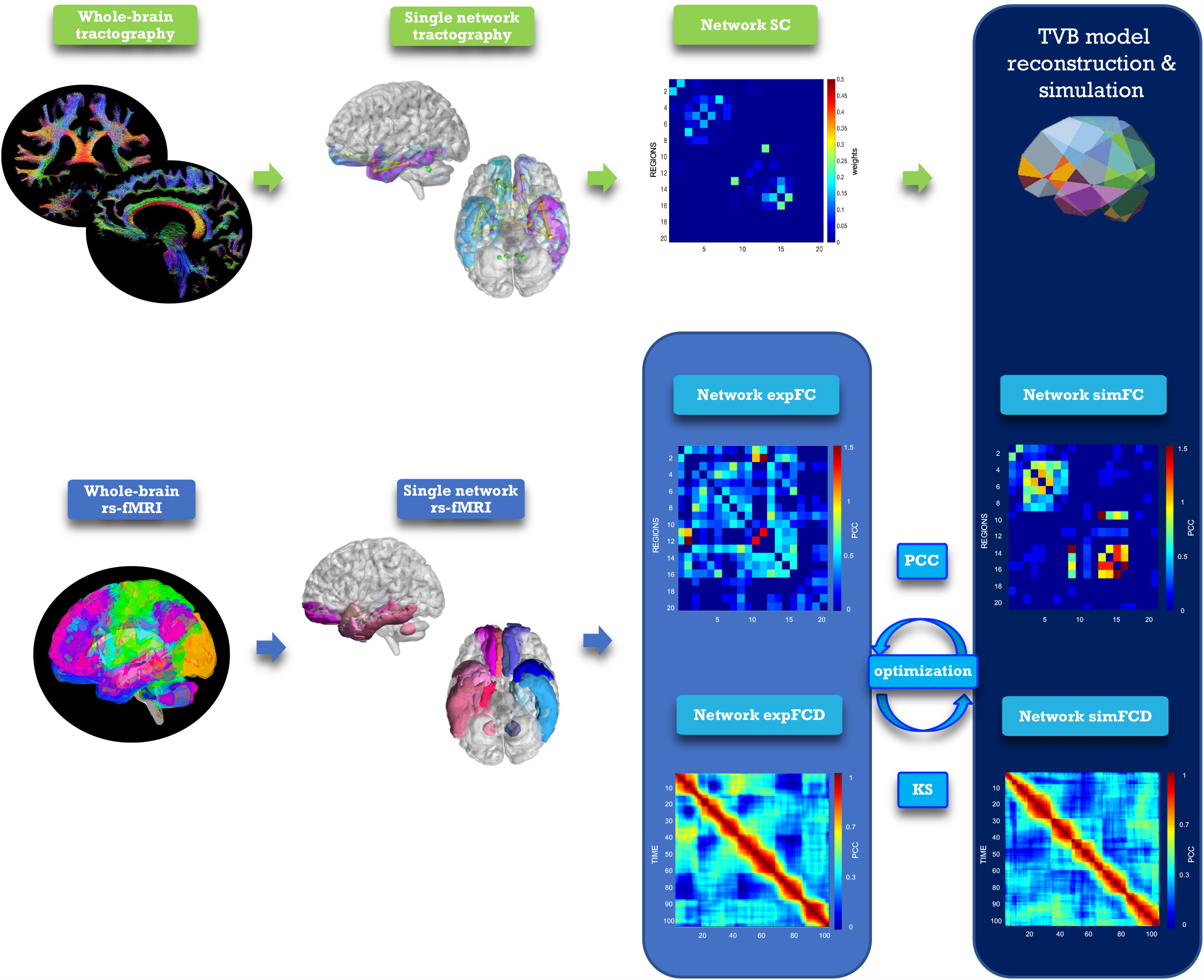
Analysis and modelling workflow. Schematic representation of MRI processing steps integrated in the modelling workflow. From top left, clockwise: diffusion-weighted images after preprocessing and tractography, extraction of a network, structural connectivity (SC) matrix reconstruction for the selected network, TVB simulation performed for the network, reconstruction of simulated static and dynamic (simFC and simFCD) functional connectivity matrices of the same network, optimization of the simulation using model inversion with the experimental FC and FCD (expFC and expFCD), derived from BOLD signals of nodes belonging to the network, as target. Optimal TVB simulation implies both the highest Pearson Correlation Coefficient (PCC) for static functional data and the lowest Kolmogorov-Smirnov (KS) distance for dynamic functional data.

### Statistical analysis

Statistical tests were performed using SPSS software version 21. Optimal TVB parameters derived for each subject and for each network were tested for normality (Shapiro-Wilk) and then two control tests were performed to assess: i) whether different networks presented a different E/I balance within the same clinical group (i.e., evaluation of the inter-network E/I balance); ii) whether inter-networks E/I balance changed in healthy vs pathological subjects. Two statistical tests were used: i) univariate general linear model followed by bias-corrected accelerated Bootstrap [37] to correct for age and gender differences in the groups and take into account non-Gaussian data distributions; ii) multivariate general linear model between the mean difference (i.e. the difference between the mean value) of TVB parameters in each network compared to the other networks in different clinical groups. Then, a multiple regression analysis was performed to investigate the relationship between individual scores of the 5 cognitive domains (memory, language-fluency, visuo-constructional abilities, attention, and executive functions) and the optimal TVB parameters. Neuropsychological scores in each cognitive domain were considered as dependent variables while model parameters derived for each network were used as predictors in a backward approach. The regression algorithm automatically removed one or more predictors to identify which of them significantly (p<0.05) explained neuropsychological scores variance.

Meaningful TVB parameters were given as an input to clustering analysis. To avoid overfitting in the study design, the clustering algorithm first performed a feature selection reducing the number of TVB parameters i) through a semi-supervised approach using LASSO regression model with TVB parameters as independent variables and the diagnostic class as dependent variable; ii) via PCC between the survived TVB parameters and the diagnostic class; iii) through Variant Inflaction Factors to find out just three meaningful but not correlated features. Then the number of clusters was derived using Gap statistics and the K-means algorithm was applied to label each subject into one cluster defining a personalized fingerprint [38].

### Code and data accessibility

All codes used for this study are open source. The optimized TVB C code can be found at https://github.com/BrainModes/fast_tvb. The dataset will be made available on Zenodo.

## RESULTS

### Networks E/I balance

Model optimization was performed in each of the six brain networks considered in this work. Global coupling (G) and mesoscopic network parameters (J_i_, J_NMDA_, w_+_) were adjusted iteratively to fit the experimental data. The reliability of the procedure was assessed by an extensive exploration of the parameter space and by iterating the optimization using different initial conditions (Suppl Fig.2) [35]. Model optimization yielded subject-specific sets of model parameters describing connectivity and E/I balance in each network. TVB parameters revealed differences between networks of healthy and pathological subjects (Suppl Fig.3; Suppl Table 2) that will be further analyzed and explained below.

### Pathology and inter-networks E/I balance

The mean difference of each network compared to the others was computed in different clinical groups for all the TVB parameters (i.e., G, J_i_, J_NMDA_, w_+_). Significant mean difference changes were found both for the TVB parameters in several networks (Fig.3) with network changes summarized in Fig.4. In particular, both in AD and FTD, the connectivity strength (G) decreased in LN and increased in DMN compared to HC; in FTD, G of FPN was lower with respect to other networks. Considering mesoscale synaptic parameters, both FTD and AD showed lower excitatory coupling (J_NMDA_) in SMN compared to HC; in FTD, J_NMDA_ was lower in VN and higher in FPN; in AD, J_NMDA_ in DMN was higher with respect to other networks. Both in AD and FTD, recurrent excitation (w_+_) increased in SMN compared to HC; in FTD, w_+_ was lower in FPN; in AD w_+_ was lower in DMN with respect to other networks. In FTD, inhibitory coupling (J_i_) was lower in FPN and higher in DMN; in AD, AN showed higher J_i_ and LN lower J_i_ with respect to other networks.

**Figure 3.**
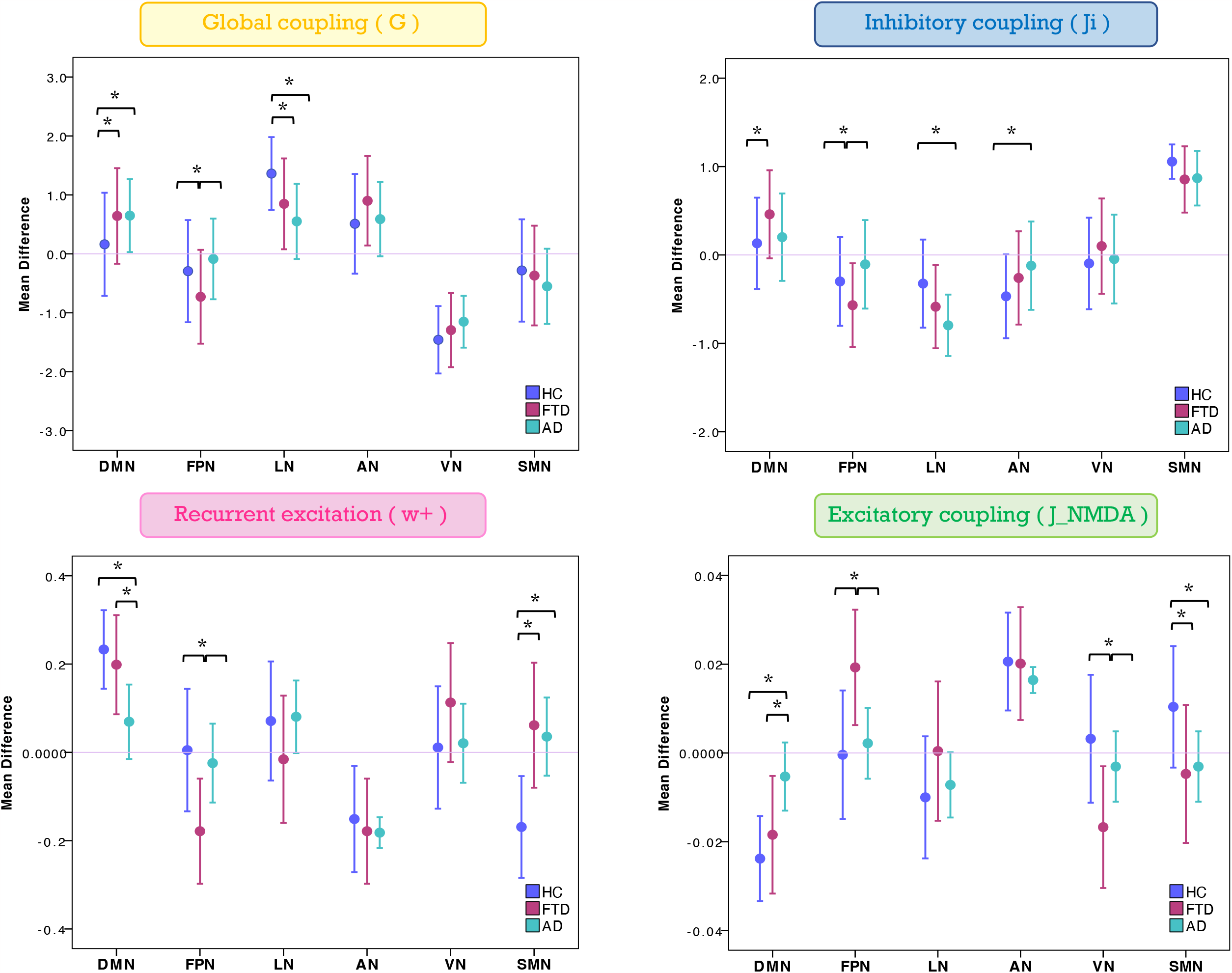
Changes of inter-network relationship. Mean difference of TVB parameters in each given network (DMN, FPN, LN, AN, VN, SMN) against the others. Positive/negative values indicate a higher/lower TVB parameter mean in a network (on the x axis) with respect to the TVB parameter mean in the others (line at mean difference 0). Asterisks indicate significant differences (p<0.05) between clinical groups (HC, FTD, AD).

**Figure 4.**
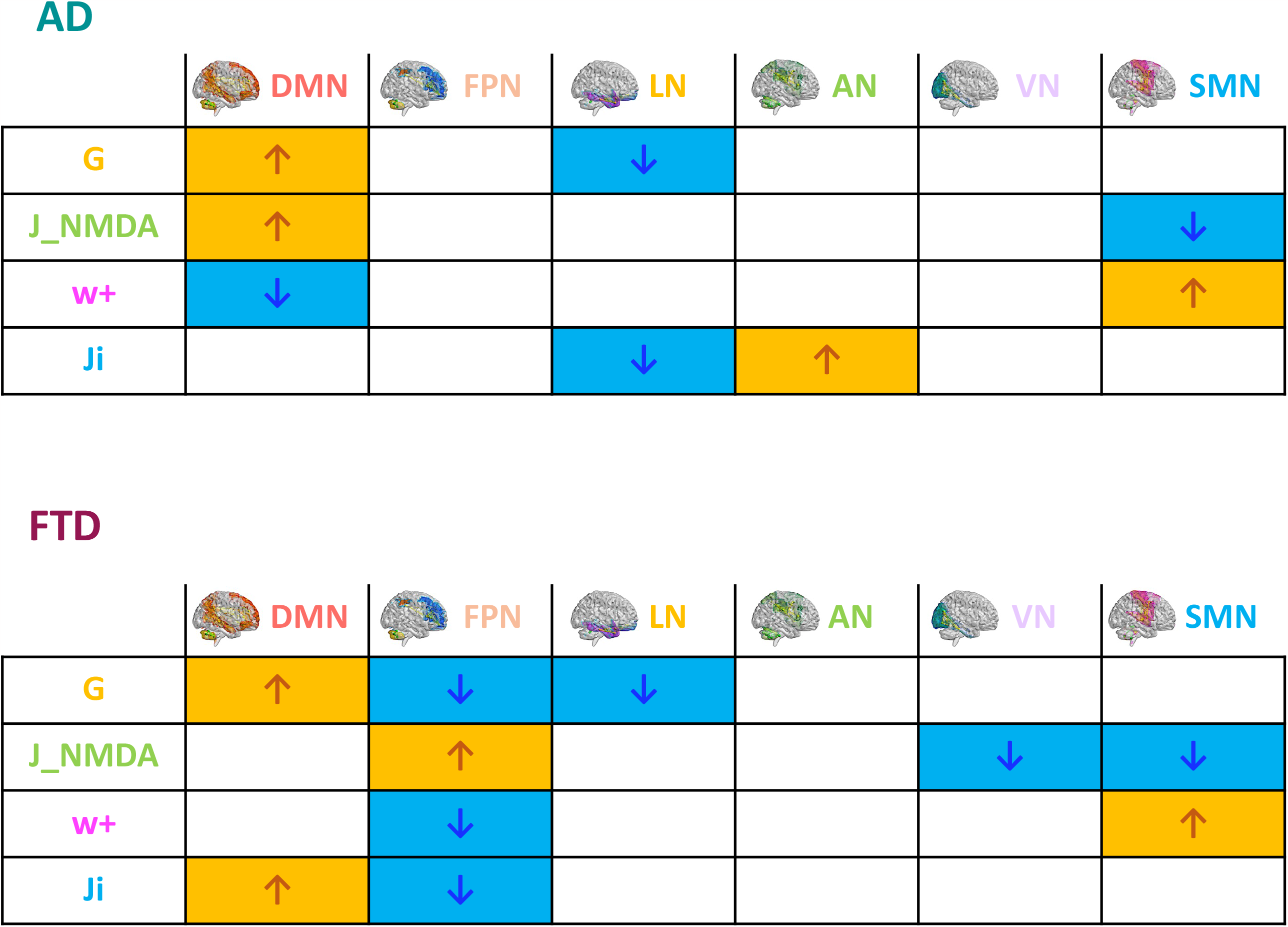
Pathological impact on inter-network relationships. Inter-network (DMN, FPN, LN, AN, VN, SMN) relationship patterns related to neurodegeneration are summarized in the tables. The increase (yellow) or decrease (blue) of network TVB parameters (G=global coupling, J_NMDA=excitatory coupling, w+= recurrent synaptic excitation, Ji= inhibitory coupling) is indicated with colored arrows.

### Clinical relevance of TVB parameters

To assess the significance of the observed mean difference changes in TVB parameters, these were used in backward regression to explain the variation of scores associated to different neuropsychological domains assessed in patients. Network-specific levels of global coupling, excitatory coupling, inhibitory coupling and recurrent excitation (predictors) significantly (p<0.05) explained a percentage of variance in the cognitive domains, in which the network is involved (Table 1). The explained variance ranged from ∼20% to ∼45%. Therefore, the mean difference changes in TVB parameters were relevant to explain the neuropsychological performance of patients.

**Table 1:** Backward regressions results.

| NETWORKS | VARIABLE<br>(Neuropsychology) | PREDICTORS<br>(TVB-parameters) | EXPLAINED<br>VARIANCE | SIGNIFICANCE |
| --- | --- | --- | --- | --- |
| VISUAL | MEMORY | J_NMDA | 21.3% | 0.027 |
|  | LANGUAGE-FLUENCY | w+, Ji | 30.1% | 0.028 |
| SOMATOMOTOR | VISUO-CONSTRUCTIONAL | Ji | 21.3% | 0.030 |
| ATTENTION | MEMORY | w+, J_NMDA, Ji | 33.4% | 0.047 |
| LIMBIC | MEMORY | Ji | 21.5% | 0.026 |
|  | ATTENTION | w+, G, J_NMDA | 42.0% | 0.030 |
| FRONTOPARIETAL | VISUO-CONSTRUCTIONAL | w+, G, J_NMDA, Ji | 45.7% | 0.027 |
|  | EXECUTIVE FUNCTION | J_NMDA | 21.3% | 0.027 |
| DMN | LANGUAGE-FLUENCY | G, J_NMDA, Ji | 39.1% | 0.022 |

The variance explained by the parameters used in backward regressions is calculated with the $R^2$ index. Significant threshold is set at $p < 0.05$ . For each cognitive domain a different combination of features significantly explains a percentage of the variance (ANOVA).

### Patients’ labelling according to network properties

The TVB parameters that significantly explained the neuropsychological performance were considered for patients’ labelling using machine learning strategies. From the nineteen parameters identified with backward regression (Table 1) the LASSO algorithm allowed to reduce them to six. Then, G of FPN was excluded, presenting PCC<0.1, and after Variant Inflation Factors three independent and not correlated variables were considered as the most informative features to perform patient’s labelling: J_i_ of AN, G of the LN and G of the DMN. Gap statistics identified that seven homogeneous classes would be appropriate and the K-means assigned each subject to one of the seven clusters. Each of the identified clusters was characterized by a specific composition of TVB network features (Fig.5A; Suppl Fig 4). Considering the biophysical meaning of each parameter, they could be described as follows:

**Figure 5.**
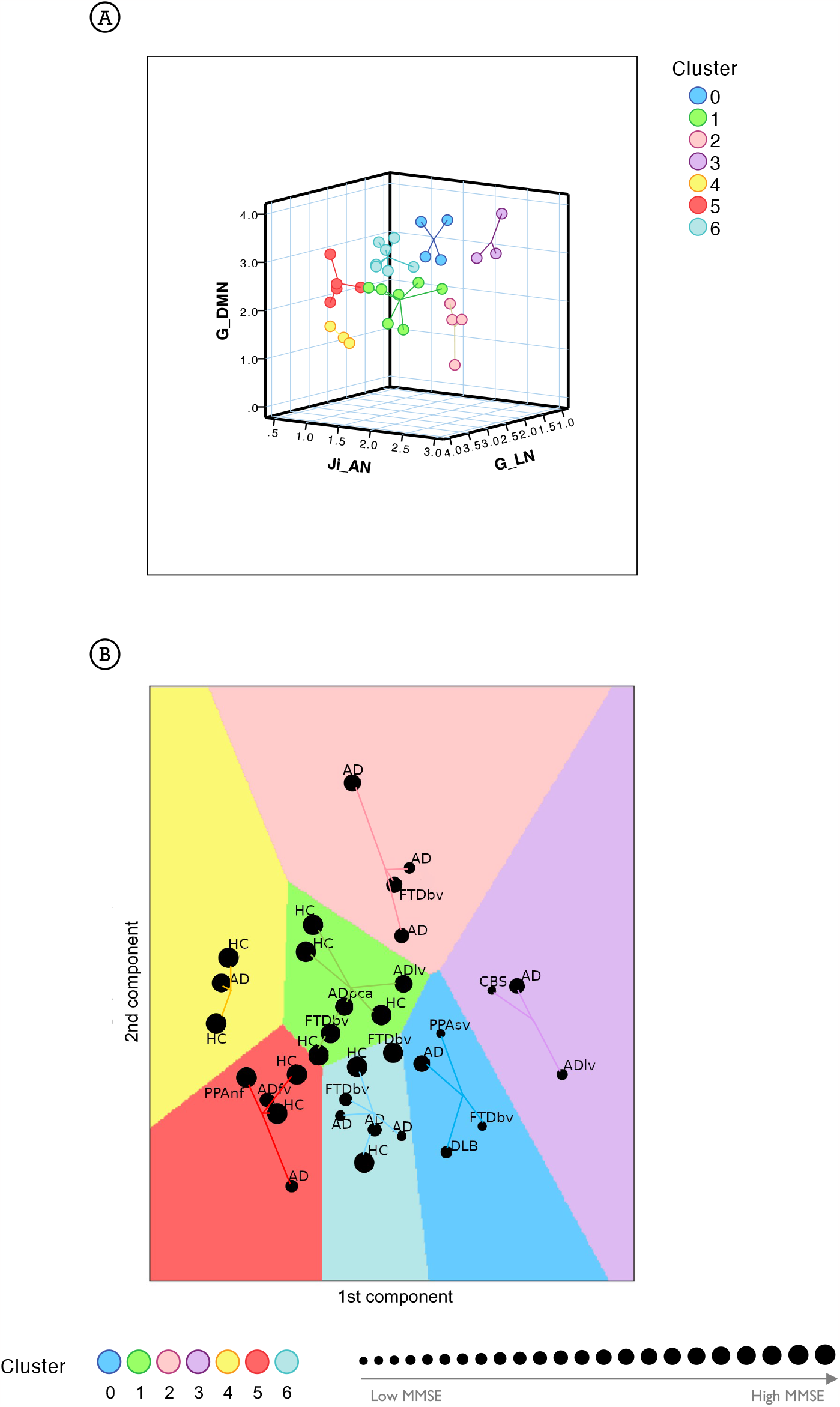
Clustering analysis. **A)** Visual representation of the seven clusters (in different colors) identified with k-means analysis using the most meaningful TVB biophysical parameters as input variables. Cognitive network properties (Ji in AN, G in LN, G in DMN) were considered as the most informative features to perform patient labelling and each of the identified clusters was characterized by a combination of low and high TVB-derived optimal parameters. Each dot represents a subject and lines connect subjects to their own cluster centroid. **B)** Each subject was assigned to one of the seven clusters (HC=healthy control, AD=typical Alzheimer’s Disease, ADlv= AD logopenic variant, ADfv= AD frontal variant, ADpca= AD posterior cortical atrophy, CBS= corticobasal syndrome, DLB= Dementia with Lewy bodies, FTD=Frontotemporal Dementia, FTDbv= FTD behavioral, PPAnf= Primary Progressive Aphasia non-fluent variant, PPAsv= Primary Progressive Aphasia semantic variant) identifying a personalized fingerprint based on cognitive network properties. Each dot represents a subject and lines connect subjects to their own cluster centroid. The dot dimension corresponds to the MMSE value.

- Cluster 0 and cluster 3 were mainly characterized by low connectivity strength of LN, high connectivity strength of DMN and hyperinhibition in AN;
- Cluster 1 and cluster 4 were mainly characterized by high connectivity strength of LN, low connectivity strength of DMN and low inhibition in AN;
- Cluster 5 and cluster 6 were mainly characterized by high connectivity strength of LN, high connectivity strength of DMN and low inhibition in AN;
- Cluster 2 was mainly characterized by low connectivity of LN, low connectivity strength of DMN and hyperinhibition in AN.

Clusters 0 and 3 were associated with the lowest mean MMSE values (20.39±5.21 and 18.57±8.28 respectively) while clusters 1 and 4 were associated with the highest mean values (29.08±1.14 and 29.33±1.16 respectively) (Fig.5B; Table 2). No HC was classified into clusters 0 or 3. Moreover, different disease phenotypes were distributed amongst the clusters (Fig.5B): typical AD subjects spread through clusters supporting a heterogeneous distribution of connectivity values in the LN and DMN networks and inhibition of the AN, but no AD patient was found in cluster 1 and the single AD patient belonging to cluster 4 presented a high MMSE score; on the other hand, cluster 0 contained the DLB phenotype, cluster 1 both the non-amnesic variants of AD (ADlv and ADpca), cluster 3 the logopenic variant and the CBS characterized by low MMSE values and cluster 5 contained the frontal variant. Considering the FTD group, FTDbv were heterogeneous and distributed amongst different clusters, but no FTDbv were found in cluster 3. On the other hand, cluster 0 contained PPAsv and cluster 5 PPAnf. Finally, pharmacological assessment of subjects belonging to different groups indicated that the majority of subjects following an antidepressant or anxiolytic treatment belonged to cluster 0 or 1 (Table 2). In particular, subjects belonging to cluster 0 were following an antidepressant therapy mainly with selective serotonin reuptake inhibitors (SSRIs), with the exception of one patient, treated with vortioxetine. Patients belonging to cluster 1, instead, were taking antidepressant drugs different from SSRIs, such as tricyclic antidepressants (e.g., amitriptyline) and serotonin-norepinephrine reuptake inhibitors (e.g. duloxetine), apart from one HC belonging to this group who was found to be on a SSRIs treatment.

**Table 2:**
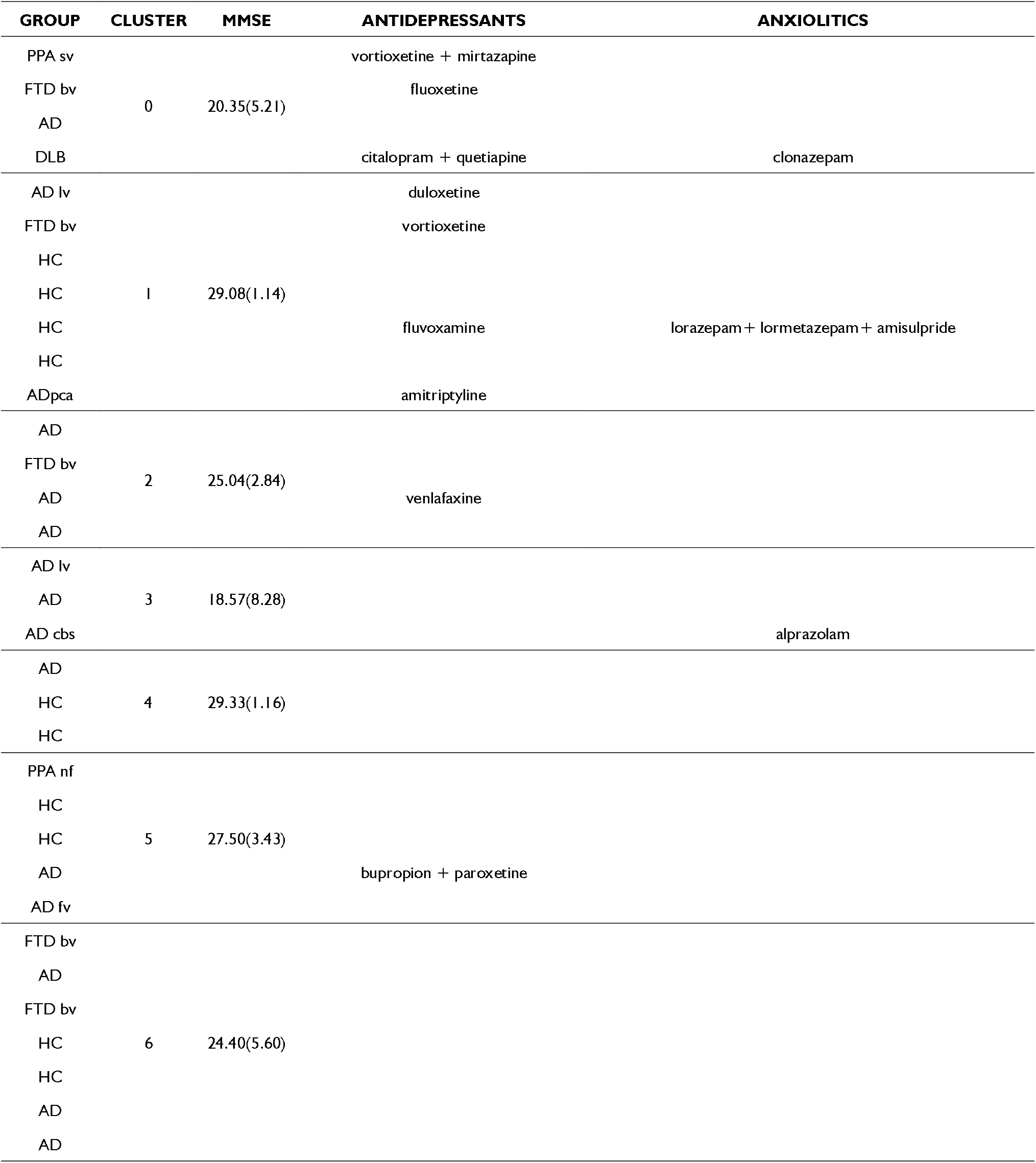
MMSE (mean,SDs) and ongoing pharmacological treatment.

## DISCUSSION

In this work we have generated virtual brain models of dementia patients and simulated neural dynamics of brain networks. The main result is the emergence of specific patterns of alteration in DMN, FPN and LN, which allow to differentiate AD from FTD. Inter-subject differences, matching the individual neuropsychological profiles and pharmacological treatment, suggest that this approach can generate personalized fingerprints of the disease that could be used to set up future stratification and interventional strategies.

### Average model parameters in brain networks of AD and FTD

In a first analysis, we compared AD and FTD for their average network model parameters. Model parameters markedly differentiated the mechanisms underlying brain networks dynamics in AD and FTD, with the most typical changes being concentrated in the DMN and LN of AD and in the FPN of FTD.

#### Integrative networks

##### Global coupling

In both pathologies, G increased in DMN and decreased in LN, while it decreased in FPN in FTD only. It is worth noting that, in these simulations, G represents the overall strength of connections between nodes inside a specific brain network. Moreover, G derives from dynamic TVB analysis and not from functional analysis on fMRI data [39], providing new insights into brain connectivity that do not necessarily compare to previously reported connectivity alterations.

In late onset AD there is meta-analytic evidence for a progressive decline of DMN FC, in particular in the posterior component (precuneus, posterior cingulate cortex)[40]. Increased FC between the posterior DMN and high connectivity hubs, mainly located in the frontal lobes, has been reported in the prodromal stages [40]. The present observation of increased G in DMN reflects hyper synchronicity, a state in which complexity is reduced along with mutual information transfer among the nodes [41]. This concept, deriving from dynamic system theory, is clearly at odd with the common belief that stronger connectivity might represent compensation, leading to the conclusion that a phase-locked hypersynchronous network can perform very limited computations [3,39]. Consistent with this hypothesis is the finding of diffused increase of spectral power in the EEG delta band of AD patients [42].

Decreased FC inside LN and from LN nodes to neighboring regions has been associated with deterioration of memory and emotional functions [43]. In FTDbv, a functional disconnection between frontal and limbic areas and an increased FC between DMN regions have been proposed as the probable correlates of apathy and stereotypic behavior [44,45]. The decreased G within LN and FPN may be also very detrimental, leading to a reduction of computational states [12,39].

#### Synaptic parameters

Another typical pattern differentiating AD from FTD emerged from synaptic parameters. Akin with neuropathology, the major AD changes were detected in DMN, while FTD changes mainly occurred in FPN. DMN showed increased excitatory coupling (J_NMDA_) and reduced recurrent excitation (w_+_) in AD, while it showed increased inhibitory coupling (J_i_) in FTD. FPN showed no changes at all in AD but it showed a complex set of changes in FTD, including increased J_NMDA_, reduced w_+_ and reduced J_i_. LN showed reduced J_i_ in AD. Therefore, the E/I balance, which remarkably impacts on brain dynamics [7], was altered in different brain networks, further differentiating AD and FTD.

We can just speculate about the meaning of these changes since information on synaptic parameters in AD and FTD pathologies is sparse. The increased J_NMDA_ in DMN may support the hyperexcitability supposed to explain cognitive impairment in AD [46]. Local hyperexcitability in the DMN was observed in previous studies, despite a net decrease in inhibitory and excitatory synaptic proteins [47,48]. The reduced J_i_ of the LN may support the limbic disinhibition reported in AD, which has been associated with a loss of GABAergic receptors [49]. The reduced J_i_ of the FPN is consistent with the reduction of GABA concentration reported in FTD, which has been associated with behavioral disinhibition [50]. Our simulations also predict overinhibition in the DMN of FTD, which provides a further differentiation with AD, where inhibition is not changed while excitation is enhanced. DMN has recently been suggested to take part in FTD pathophysiology [51]. Therefore, the patterns of synaptic changes captured by our study prompts for further experimental and model analysis of synaptic alterations in microcircuits of the AD and FTD brain.

#### Motor and sensory networks

Both in AD and FTD, the SMN showed reduced J_NMDA_ and increased w_+_. Although the impairment of GABAergic and glutamatergic systems in the motor and sensory networks still needs to be clarified, it should be noted that motor dysfunctions are known to occur in both AD and FTD [52,53]. In AD, a reduced motor cortex excitability has been reported in mild cognitive impairment [55], suggesting that these parameters may change along the evolution of the disease. In FTD, motor circuit abnormalities have been suggested to depend on altered glutamatergic transmission [54]. Interestingly, in FTD abnormalities of oculomotor functions have been reported [56], which might be linked not only to SMN impairment, but also to a more extended involvement of VN, as supported by our results.

### The relationship between network neurophysiology and neuropsychology

Model parameters for individual subjects were correlated with behavioral observations. Global coupling and synaptic parameters of each network significantly contributed to explain neuropsychological scores in specific cognitive domains: LN, AN and VN with memory; DMN and VN with language-fluency; LN with attention; SMN and FPN with visuo-constructional performance; FPN with executive functions. This evidence is in line with several reports on the importance of motor regions in visuo-constructional performance [57], the contribution of AN and limbic areas in memory [58], the relevance of frontoparietal areas for executive and visuo-constructional control [59,60], the role of DMN integration for semantic fluency [61], and the involvement of visual structures in memory and language-fluency [62,63].

Thus, the relationship between neurophysiological parameters in brain networks and neuropsychological scores, which has not been investigated before, supports the clinical relevance of model parameters and provides important cues for understanding the physiopathology of AD and FTD.

### Toward personalized fingerprints of AD and FTD patients

The most meaningful model biomarkers for patient’s labelling were G in DMN, G in LN, J_i_ in AN, consistent with known salient aspects of dementia affecting the ability of daydreaming (DMN), emotional control (LN) and attention (AN). Subjects were found to be distributed between seven different clusters revealing correspondence with their cognitive status (assessed with MMSE) and pharmacological treament.

#### Cognitive performance

Patients with the lowest MMSE belong to clusters with low G in LN, high G in DMN and high J_i_ in AN (clusters 0, 2, 3 in Fig.5), while HC are not found in these clusters at all. Patients with relatively high MMSE, as well as HC (MMSE>30), spread over clusters with high G in LN, low G in DMN and low Ji in AN (clusters 1, 4, 5, 6 in Fig.5). These observations highlight the importance of DMN, LN, and AN connectivity strength and E/I balance to ensure healthy cognitive function. Interestingly, high G between DMN nodes is associated with a worse performance, being hence disruptive and not compensatory.

Our findings suggest that the heterogeneity of subject-specific TVB parameters is able to identify AD “subtypes” [64,65] and FTD variants. Indeed, subjects belonging to atypical forms of AD and FTD variants were assigned to different clusters, capturing specific aspects of these pathologies and mostly mapping clinical severity assessed with MMSE. A finer grained analysis based on clinical phenotypes is not currently possible, given the limited sample size.

#### Pharmacological treatment

Patients’ labelling based on TVB parameters was found to correlate with pharmacological treatment. Most subjects belonging to clusters 0 and 1 were on antidepressant or anxiolytic treatment (cf. Table 2), which suggest that they may influence the connectivity strength and the E/I balance of cognitive networks. The effect of SSRIs on LN and DMN FC is increasingly recognized in literature [66,67]. The effect of antidepressant treatment with molecules different from SSRIs, such as vortioxetine, tricyclic molecules or SNRIs [68], as well as the influence of antidepressants on GABA and glutamate levels needs further assessment [69]. Considering that patients treated with SSRIs belong to cluster 0 while patients treated with other antidepressant classes belong to cluster 1, our results pose a very intriguing question: is there an opposite impact on cognitive networks exerted by antidepressants with different mechanisms of action or does the cognitive networks profile determine pharmacological treatment response? Future work should study TVB parameters longitudinally pre-post treatment to answer this important question with major potential clinical impact.

It should be noted that, in our sample, patients were not treated with NMDA receptor antagonists (like memantine) [87] or acetylcholinesterase inhibitors (like galantamine, rivastigmine, and donepezil) [71], which are also known to act on AD. NMDA receptors are main triggers of synaptic plasticity and excitotoxicity [72] and cholinergic receptors also impact on synaptic plasticity and learning [73]. Since J_NMDA_ in the Wong-Wang neural mass model is mostly related to slow synaptic mechanisms driven by NMDA receptors [7] and receptor density can be remapped onto TVB [74], an assessment of these receptor-dependent functions could be an important development in future studies.

## STUDY CONSIDERATIONS

The small sample size can be seen as a potential limitation in the present study. However, the main aim of this investigation was to assess the ability of TVB to provide a personalized fingerprint of patients, potentially beyond known diagnosis. Virtual brain modelling provides a set of features at single subject level, otherwise not available from standard signal/image analysis, which can be grouped thereafter using appropriate statistical analysis. Thus, the small sample size does not impact significantly on the TVB capacity of uncovering subject-specific features of connectivity, and E/I profile. The high correlation of TVB parameters with both cognitive performance and pharmacological treatment reveals indeed its exquisite sensitivity to single-subject features and opens a broad perspective for clinical applications. On the other hand, the application of TVB to a larger cohort bears the prospect of improving disease classification of subtypes and of establishing interventional workflows.

## CONCLUSIONS AND PERSPECTIVES

The present study demonstrates that brain networks can be characterized in terms of a meaningful set of mesoscale parameters of network connectivity strength and E/I balance at the single-subject level in humans *in vivo*. These parameters are obtained by embedding structural and functional MRI data into a virtual brain model (TVB). TVB parameters measured in AD and FTD yield subject-specific profiles correlated with neuropsychological, pharmacological and clinical scores. At present, it is unclear whether network properties influence or are influenced by therapy suggesting that future studies should systematically address this issue to effectively instruct therapeutic choices. The identification of network abnormalities in patients may be used to design neuromodulation, neuropharmacological and neuropsychological paradigms capable of regulating circuit function and plasticity [75]. In aggregate, TVB parameters are shedding light on the changes occurring inside the brain networks of AD and FTD patients opening new perspectives for understanding disease mechanisms and for designing personalized therapeutic approaches. Future studies should involve larger cohorts of patients to confirm the implications of this first investigation for dementia assessment and management.

## Abbreviations

expFC: experimental Functional Connectivity
expFCD: experimental Dynamic Functional Connectivity
FC: Functional Connectivity
PCC: Pearson Correlation Coefficient
SC: Structural Connectivity
simFC: simulated Functional Connectivity
simFCD: simulated Dynamic Functional Connectivity
TVB: The Virtual Brain
AD: Alzheimer’s Disease
FTD: Frontotemporal Dementia

## AUTHORS CONTRIBUTION

Patients’ recruitment and clinical assessment: MCR, AC

MRI recordings: LF, AP, LM

Neuropsychological testing: FC

Data analysis: AR, AM, FP, MS

TVB modelling and simulation: AM, FP, MTM, FA, ED, CWK

MRI theory and protocol design; CWK, FP

TVB support: MS, VJ, PR

Neurological support: SC, AC, MCR

Paper writing: AM, ED, FP

Work coordination and paper finalization; ED, CWK, FP

All Authors have contributed to paper discussion and approved the final version of the paper.

