## Supplementary materials for "Virtual brain simulations reveal network-specific parameters in neurodegenerative dementias"

**SUPPLEMENTAL MATERIAL to the paper**

**Supplementary Figure 1| The Wong-Wang Model.** Each node of this dynamic mean field model contains one excitatory (E) and one inhibitory (I) neural population. Brain dynamics are described by a set of coupled non-linear stochastic differential equations, in which where r_i_^(E,I)^ denotes the firing rate of the excitatory and inhibitory populations, S_i_^(E,I)^ identifies the average excitatory or inhibitory synaptic gating variables at local area, i, and I_i_^(E,I)^ is the input current to the excitatory and inhibitory populations at local area, i. Inter-areal connections are weighted by the structural connectivity matrix and scaled by the global coupling G. The excitatory synaptic coupling (J_NMDA_) from the excitatory to the inhibitory population is NMDA mediated, as well as the local recurrent excitation (w_+_), while the feedback inhibitory coupling (J_i_) is GABAergic.

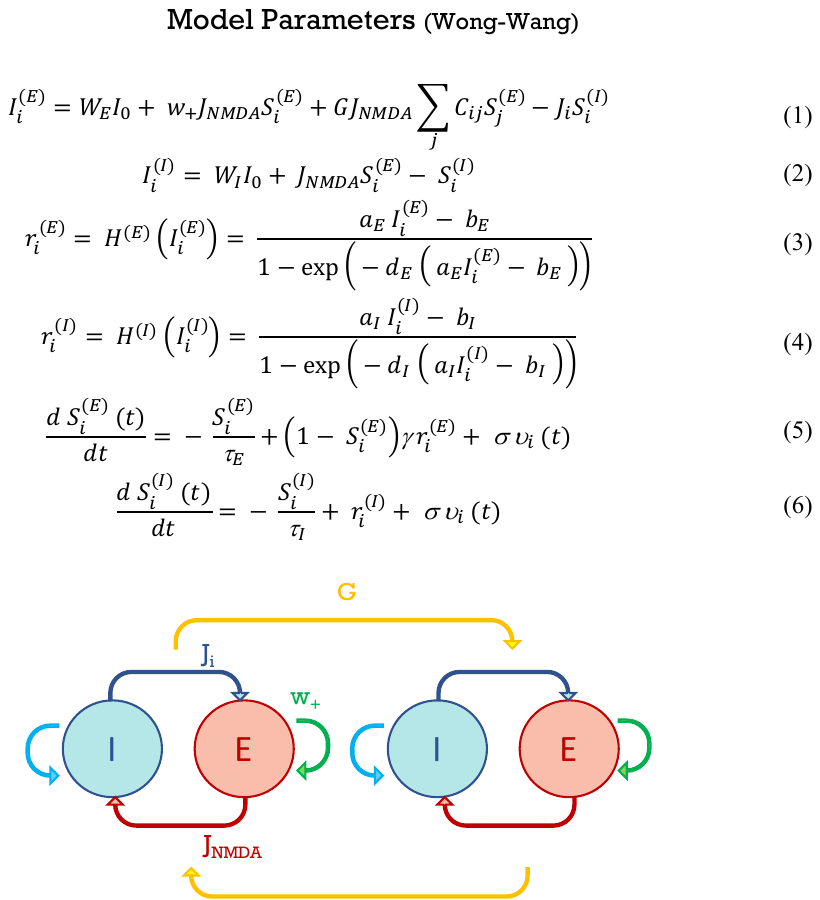

**Supplementary Figure 2| Parameter space exploration. A)** The figure shows the two steps of 2D parameter space exploration: (left) G and J_i_ are optimized first, while fixing the other two parameters (J_NMDA_ and w_+_) at their standard value; (right) G and J_i_ are then fixed at their optimized value and J_NMDA_ and w_+_ are optimized in turn. Heat maps represent the value of the cost function obtained with different parameters combinations. The lowest cost function corresponds to the best fit of simulated to experimental data (cf. Fig. 2). **B)** Example of iterations with randomized initial conditions. It should be noted that the optimal parameter estimates (red dots) fall in a narrow region across iterations. The histograms show the frequency of occurrence of optimal TVB parameter values. Note that the modal values of parameter distributions are almost coincident with the values used for TVB simulations (red asterisk).

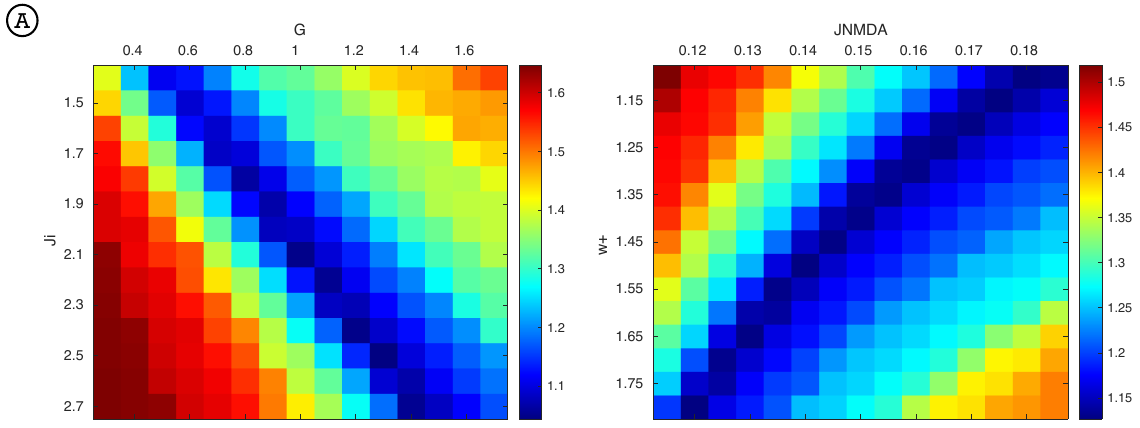

**Supplementary Figure 3|** **Boxplots of networks optimal biophysical parameters.** Boxplots of optimal TVB-parameters (G=global coupling J_NMDA=excitatory coupling w+=recurrent excitation Ji=inhibitory coupling) among networks in different clinical groups (HC=Healthy Controls AD=Alzheimer’s Disease FTD=Frontotemporal Dementia). Asterisks indicate significant differences between all the networks which are reported in detail in Supplementary Table 2 (univariate general linear model followed by bias-corrected accelerated Bootstrap, p<0.05). Each network is characterized by its own specific excitatory/inhibitory balance.

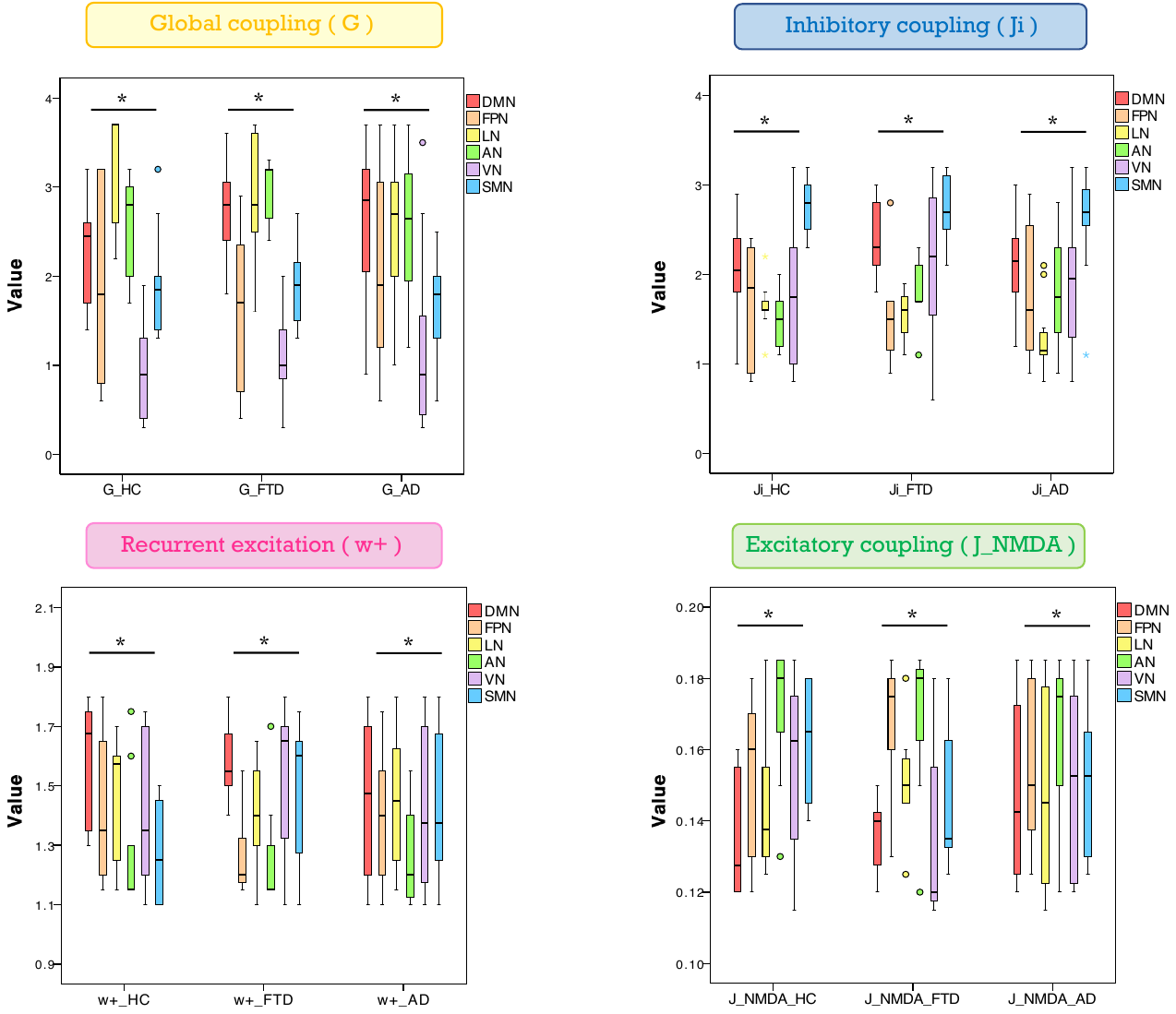

**Supplementary Figure 4| Clusters profile.** Each cluster is characterized by a typical profile of cognitive networks properties (the inhibitory coupling of attention network, the global coupling of the limbic network and the global coupling of the default mode network). The color-bar (from blue to red) represents the scale from low to high of each TVB parameter.

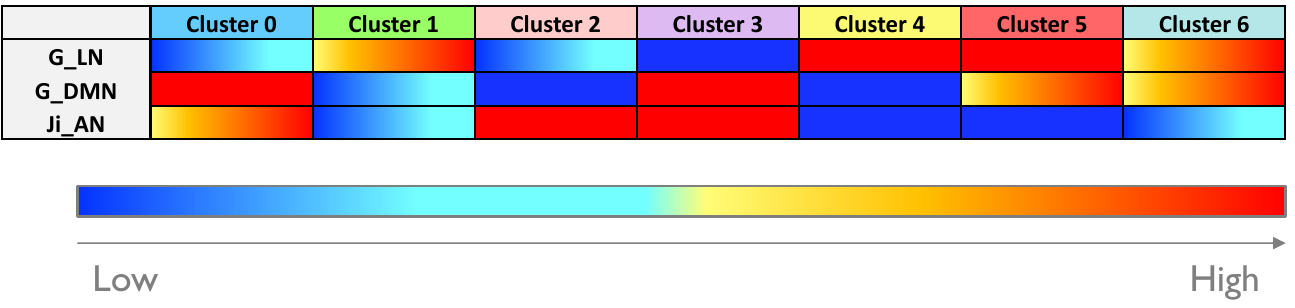

**Supplementary Table 1: Demographics, clinical and neuropsychological data.**

| **Measures** | **HC** | **AD** | **FTD** |
| --- | --- | --- | --- |
| **Males/females** | 4/6 | 3/13 | 6/1 |
| **Age (years)** | 67 ± 3 | 70 ± 8 | 69 ± 5 |
| **Education (years)** | 11 ± 5 | 9± 5 | 8 ± 4 |
| **Memory** | - | 0.6± 0.6 | 2.3 ± 1.4 |
| **Language-fluency** | - | 1.5± 1.3 | 1.6 ± 1.6 |
| **Visuo-constructional** | - | 0.9 ± 1.6 | 2.8 ± 1.7 |
| **Attention** | - | 0.2 ± 0.4 | 1.2 ± 1.5 |
| **Executive functions** | - | 1.2± 0.7 | 2.2 ± 1.0 |

Gender, age, education and neuropsychological scores are reported for each group (Healthy controls, HC, Alzheimer’s disease, AD, Frontotemporal dementia, FTD). Data are reported as mean±standard deviation.

**Supplementary Table 2| Network-dependent connectivity and excitatory/inhibitory balance.** The table reports significant differences of parameter (yellow: G=global coupling, green: J_NMDA=excitatory coupling, pink: w+=recurrent excitation, blue: Ji=inhibitory coupling) of each network with respect to others (horizontal lines: VN, SMN, AN, LN, FPN, DMN) in different clinical conditions (vertical columns: HC=healthy control, AD=Alzheimer’s disease, FTD=Frontotemporal dementia).

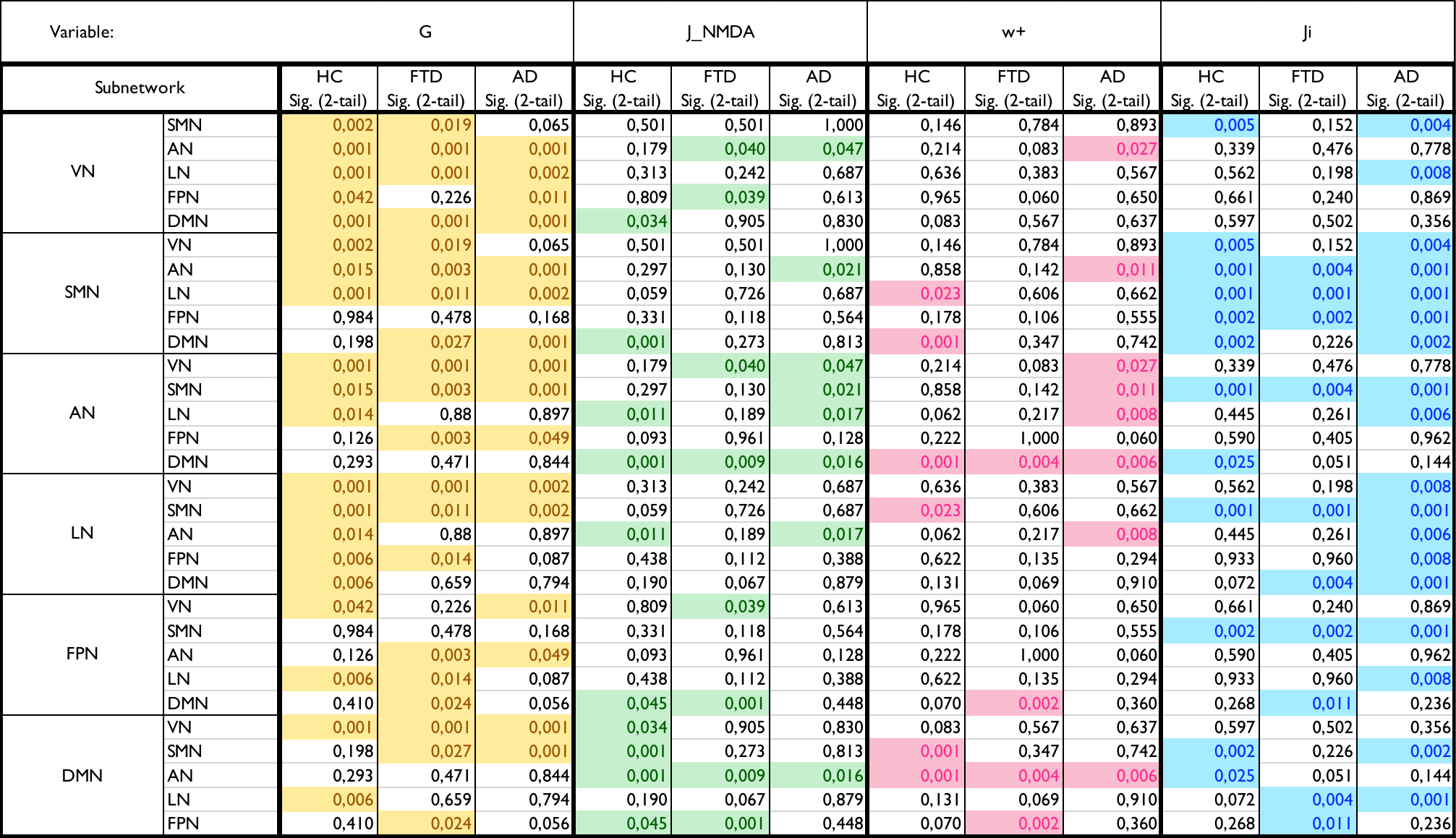
